# High quality genome and transcriptome data for two new species of *Mantamonas*, a deep-branching eukaryote clade

**DOI:** 10.1101/2023.01.20.524885

**Authors:** Jazmin Blaz, Luis Javier Galindo, Aaron A. Heiss, Harpreet Kaur, Guifré Torruella, Ashley Yang, L. Alexa Thompson, Alexander Filbert, Sally Warring, Apurva Narechania, Takashi Shiratori, Ken-ichiro Ishida, Joel B. Dacks, Purificación López-García, David Moreira, Eunsoo Kim, Laura Eme

**Author notes:** Corresponding authors: Laura Eme, Eunsoo Kim.

## Abstract

Mantamonads were long considered to represent an “orphan” lineage in the tree of eukaryotes, likely branching near the most frequently assumed position for the root of eukaryotes. Recent phylogenomic analyses have placed them as part of the “CRuMs” supergroup, along with collodictyonids and rigifilids. This supergroup appears to branch at the base of Amorphea, making it of special importance for understanding the deep evolutionary history of eukaryotes. However, the lack of representative species and complete genomic data associated with them has hampered the investigation of their biology and evolution. Here, we isolated and described two new species of mantamonads, *Mantamonas vickermani* sp. nov. and *Mantamonas sphyraenae* sp. nov., for each of which we generated transcriptomic sequence data, as well as a high-quality genome for the latter. The estimated size of the *M. sphyraenae* genome is 25 Mb; our de novo assembly appears to be highly contiguous and complete with 9,416 predicted protein-coding genes. This near-chromosome-scale genome assembly is the first described for the CRuMs supergroup.

## Background and Summary

Free-living heterotrophic flagellates play important roles in the nutrient cycling of marine and freshwater ecosystems. However, the extent of their genomic diversity is still dramatically uncharacterized. Amongst the lesser-known of these is *Mantamonas*, a genus of marine gliding flagellates initially described as very divergent from all other known eukaryotes^1^. Although *Mantamonas* was originally thought to be related to the poorly-known lineages Apusomonadida and Ancyromonadida, based on ribosomal RNA gene phylogenies and some of their morphological characteristics^1^, recent transcriptome-based phylogenomic analyses instead robustly placed *Mantamonas plastica* as sister to a clade comprising Collodictyonidae (also known as diphylleids) and Rigifilidae, altogether forming the “CRuMs” supergroup^2,3^. This clade presents diverse cell morphologies and branches at the base of Amorphea^2,4^ (Amoebozoa plus Obazoa, the latter including animals and fungi, among others). The genomic exploration of members of this supergroup therefore represents an important resource for uncovering the characteristics of this deep-branching clade, and may help us better understand evolutionary transitions within the eukaryotic tree of life, such as the acquisition of complex multicellularity in several lineages of the Obazoa. However, to date, only partial transcriptomic data is available for a handful of CRuMs taxa, including *M. plastica*^*2,3*^. Here, we isolated and described two new species of mantamonads, *Mantamonas sphyraenae* sp. nov. and *Mantamonas vickermani* sp. nov., and generated a high-quality nuclear genomic assembly for the former and transcriptomic assemblies for both species.

Overall, the cell morphology and behavior under light microscopy of these two new species (Fig. 1 and Movie S1) are comparable to what was reported in the original description of the genus *Mantamonas*^*1*^ and to our own observations of the type strain of *M. plastica*. Nonetheless, our strains appear to be slightly smaller than the 5×5 µm dimensions of *M. plastica*. Cells of this genus have one anterior and one posterior flagellum. They are flattened and somewhat plastic, with shapes ranging from wide, with more or less pointed lateral “wings” resembling the fins of a manta ray, to kite-shaped, to oval, to spherical. The left side of the cell body often displays a characteristic blunt projection, which we sometimes observed in our new strains, although less conspicuously (Fig. 1; formal species description in the Supplementary Information file).

**Fig. 1.**
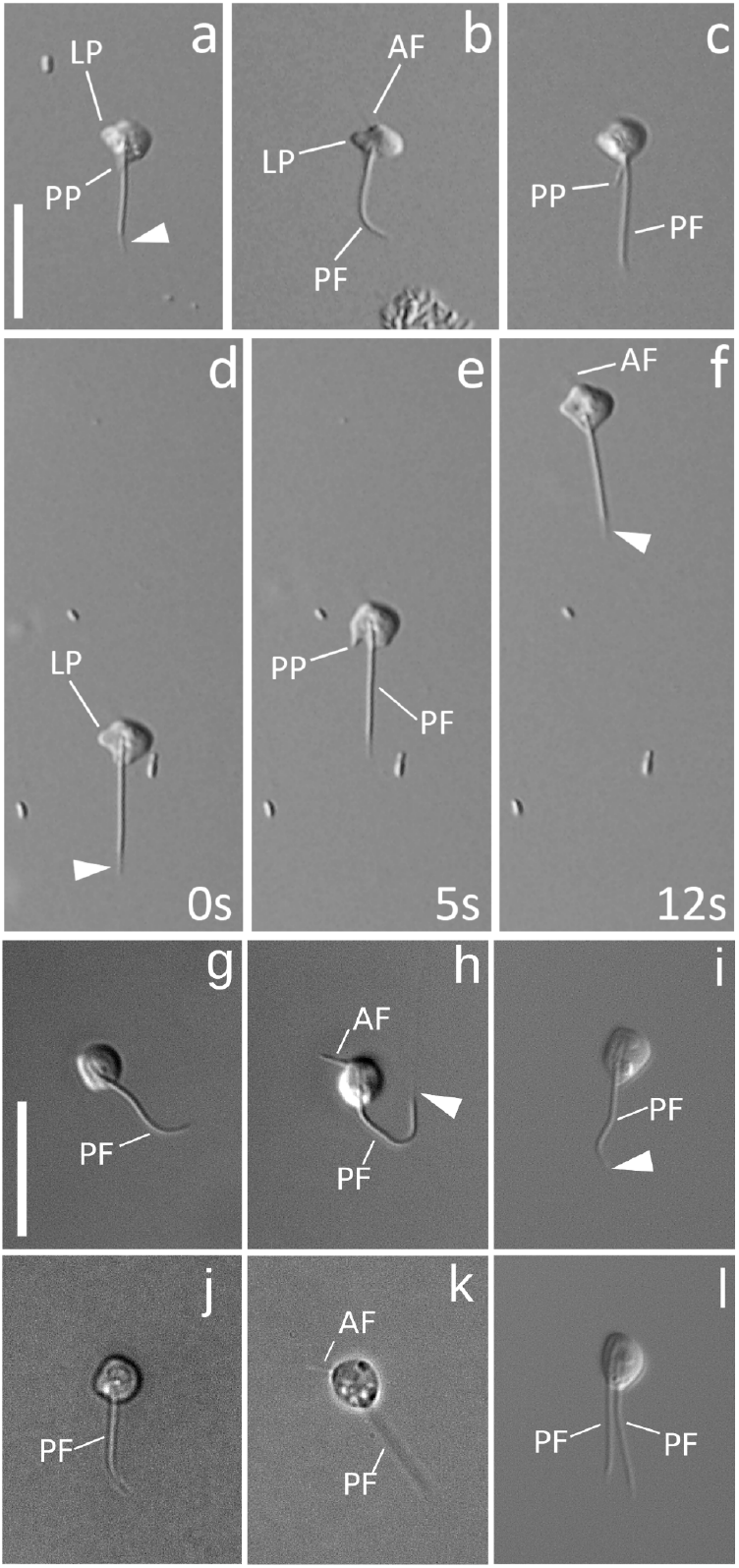
General morphology of *Mantamonas sphyraenae* sp. nov. and *Mantamonas vickermani* sp. nov. a-c) Differential interference contrast light micrographs of living *M. sphyraenae* cells. Note acroneme (white arrowheads), most visible in panel (a) but present in all micrographs. The extremely thin anterior flagellum is visible in panel (b). The left projection, present in all cells, is most distinct in (b). A posterior protrusion is often visible, usually parallel and immediately adjacent to the posterior flagellum (a,b), but sometimes at an angle to it (c). d-f) Individual *M. sphyraenae* cell imaged over a 12-second period; numbers in lower right indicate elapsed time in seconds. Note the plastic nature of the cell and lack of movement of the posterior flagellum except to trail behind the cell body. g-l) Phase and differential interference contrast light micrographs of living interphase *M. vickermani* cells. Note contrast between thick and long posterior flagellum and thin and short anterior flagellum in (g,k). l) Laterally dividing cell of *M. vickermani* with two posterior flagella. Scale bars: 10 µm. AF = anterior flagellum; LP = left projection; PF = posterior flagellum; PP = posterior protrusion; arrowhead = acroneme.

All previously known mantamonad strains were isolated from marine sediments^1^, which was also the case for our strain *M. vickermani* sp. nov., isolated from marine lagoon sediment. However, we isolated the other strain (*M. sphyraenae* sp. nov.) from the skin surface of a barracuda, which could suggest that either this species is epizootic (normally inhabiting the skin of the fish) or that the cells that we isolated were dislodged from their normal habitat and adhered to the fish skin by chance. Additional sampling and culturing efforts should help resolve this matter.

The assembled nuclear genome sequence of *M. sphyraenae* is highly contiguous (Table 1). This genome sequence was generated using long (PacBio) and short (Illumina) reads (see Methods). The average sequencing coverage was 112x for PacBio and 115x for Illumina. Three different genome assembly strategies, using Canu^5^, FALCON^6^, and MaSuRCA^7^, yielded comparable results (Supplementary Table 1), with >90% representation of the 255 Benchmarking Universal Single Copy Orthologs (BUSCO^8^) of the eukaryota_odb10 dataset (Fig. 2), indicating high completeness. For downstream analyses, we opted to use the FALCON assembly because it was the most contiguous of the three, with the majority of the contigs (59 out of 78 primary contigs) bearing telomeric repeats at both ends. In addition, 14 of the remaining contigs had telomeric repeats at one end, leading to an estimation of 66 pairs of chromosomes in the *M. sphyraenae* nucleus. Biallelic single nucleotide polymorphism (SNP) frequencies cluster around a ratio of 0.5/0.5 for each major/minor allele (Fig. 3a). This is indicative of a diploid genome, which was also supported by the statistical model of SNP frequency distribution (Supplementary Table 2).

**Table 1.** Genomic and transcriptomic assemblies statistics for *Mantamonas sphyraenae* sp. nov. and *Mantamonas vickermani* sp. nov. Values within parentheses correspond to primary plus associate contigs produced by FALCON.

|  | *Mantamonas sphyraenae* | *Mantamonas vickermani* |
| --- | --- | --- |
| Assembly type | genome | transcriptome |
| Assembly length (Mb) | 25.06 (31.49) | 21.05 |
| Number of contigs | 78 (199) | 9,668 |
| Contig mean length (Kb) | 321.30 | 2.178 |
| Longest contig (Kb) | 751.365 | 22.29 |
| Shortest contig (Kb) | 17.266 | 0.0297 |
| N50 (Kb) | 375.07 | 3.05 |
| L50 | 26 | 2035 |
| GC content | 59.19 | 46.02 |
| Total repeat content | 12.12 | -- |

**Table 2.** Summary of sequencing data records.

| Species and strain name | Type | Platform | Read type | SRA accession number |
| --- | --- | --- | --- | --- |
| *M. sphyraenae* STR306 | DNA | PacBio RS II | Single | SRR21818797 |

**Table 2.** Summary of sequencing data records.
|  |  |  | molecule |  |
| --- | --- | --- | --- | --- |
| *M. sphyraenae* STR306 | DNA | Illumina HiSeq 2500 | Paired | SRR21818798 |
| *M. sphyraenae* STR306 | DNA | Illumina HiSeq 2500 | Mate pair | SRR22188164 |
| *M. sphyraenae* STR306 | RNA | Illumina HiSeq 2500 | Paired | SRR21818794 |
| *M. vickermani* CRO19MAN | RNA | IlluminaHiSeq 2500 | Paired | SRR21818793 |

**Fig. 2.**
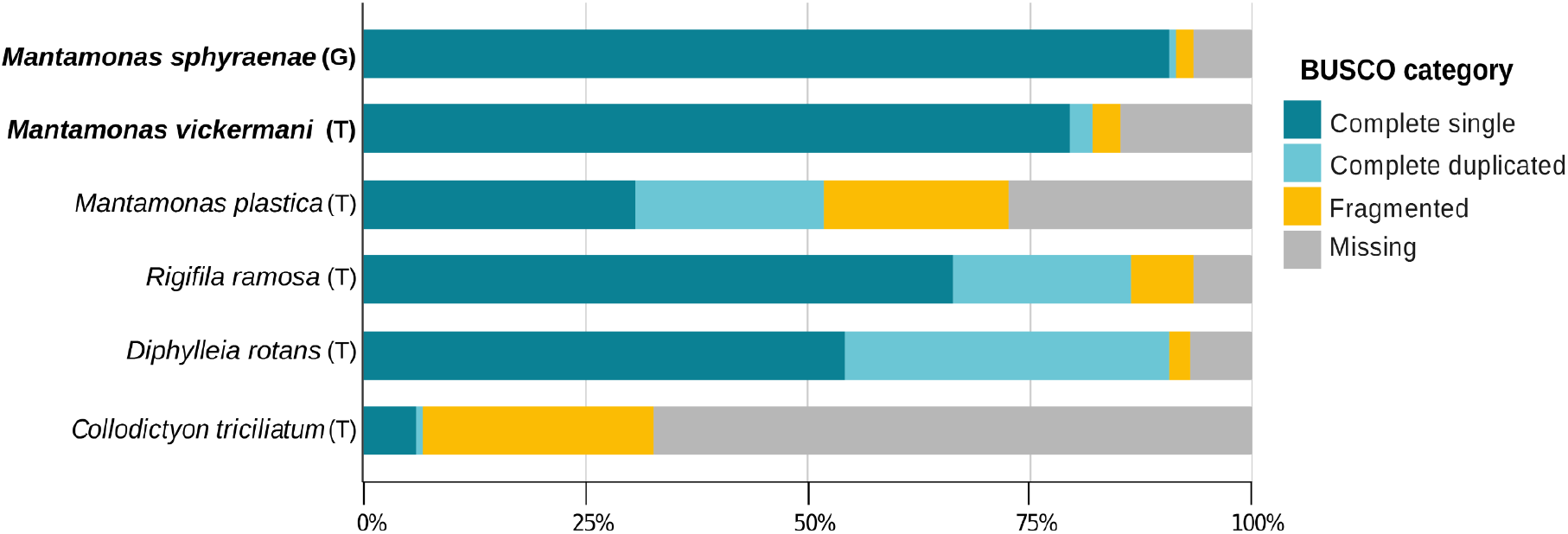
Distribution of BUSCO orthologs in inferred proteomes of mantamonad assemblies (in bold) in comparison with those of other members of the CRuMs supergroup. Proteomes were inferred from genome (G) and transcriptome (T) assemblies.

**Fig. 3.**
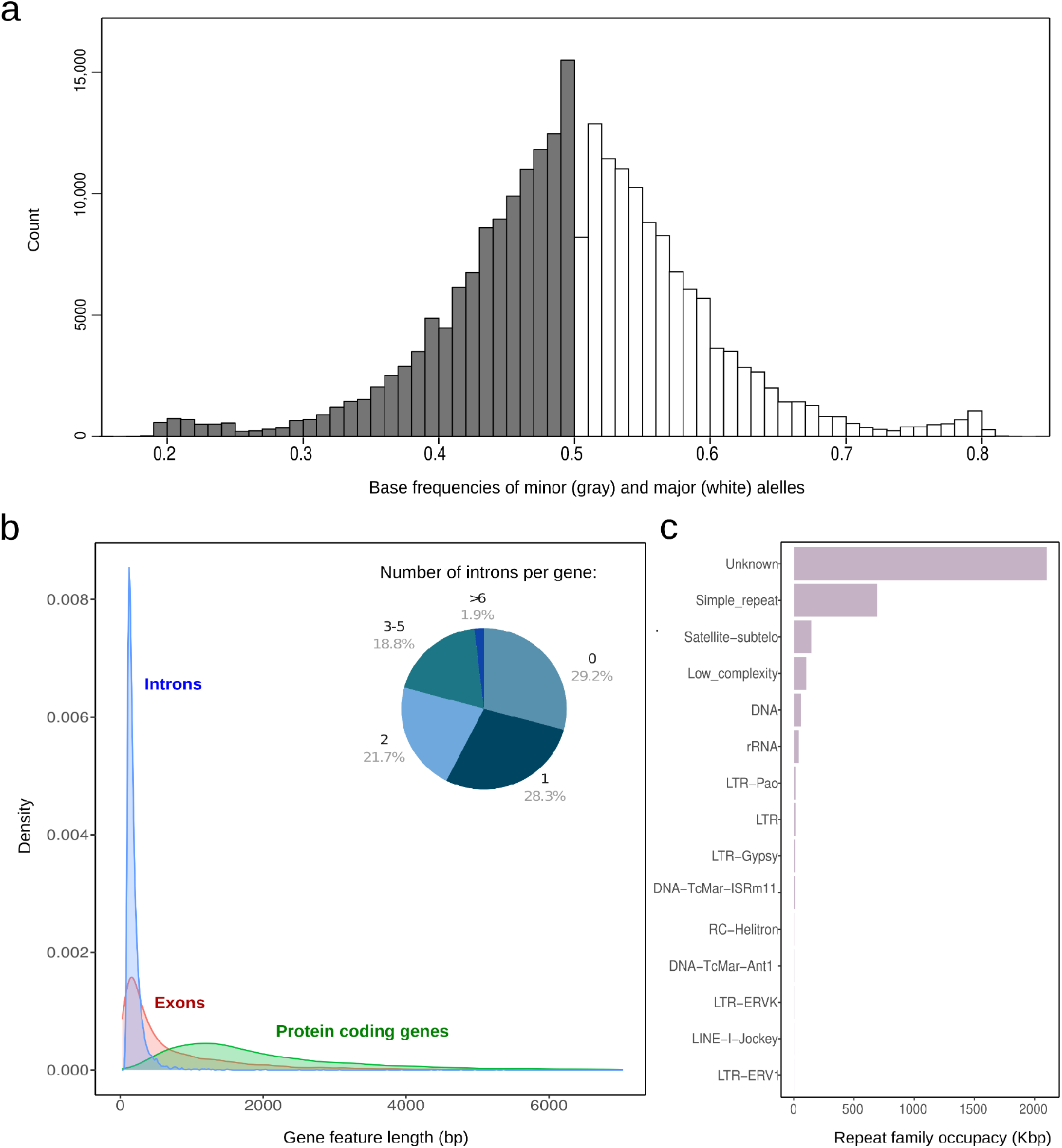
Genomic features of *Mantamonas sphyraenae* sp. nov. a) Biallelic SNP frequency distribution. b) Length distribution and intron frequency of protein-coding genes. c) Genomic occupancy of the families of repetitive elements identified *de novo*.

We predicted 9,416 protein-coding sequences in the *M. sphyraenae* genome. Genes have an average length of 2,282 bp and are mostly mono-exonic (Fig. 3b). *De novo* characterization of repetitive elements indicates that around 12% of the genome is represented by transposable elements and other repeats. While some of these were classified into different families of DNA transposons and long terminal repeat (LTR) retroelements, the vast majority comprises unclassified types (Fig. 3c).

The *de novo* assembled transcriptome of *M. vickermani* had an average sequencing coverage of 80x (Supplementary Fig. S2) and led to the inference of 9,561 non-redundant proteins. As with the genome of *M. sphyraenae*, the proteome inferred from this transcriptome resulted in a high BUSCO score, indicating the presence of a near-complete genetic complement (Fig. 2).

We inferred the phylogenetic relationships of our species within the CRuMs clade using publicly available data to reconstruct a dataset of 182 conserved protein markers. Consistent with previous studies, our maximum likelihood (ML) and Bayesian inference (BI) phylogenetic trees recovered the monophyly of CRuMs with high BI posterior probability (0.99) and ML bootstrap support (95%), although it is worth noticing that the outgroup is highly reduced since resolving the position of CRuMs in the tree of eukaryotes is outside the scope of this paper. The monophyly of *Mantamonas* received full support from both methods. We found *Mantamonas sphyraenae* to be sister to a maximally-supported clade containing *M. plastica* and *M. vickermani* (Fig. 4).

**Fig. 4.**
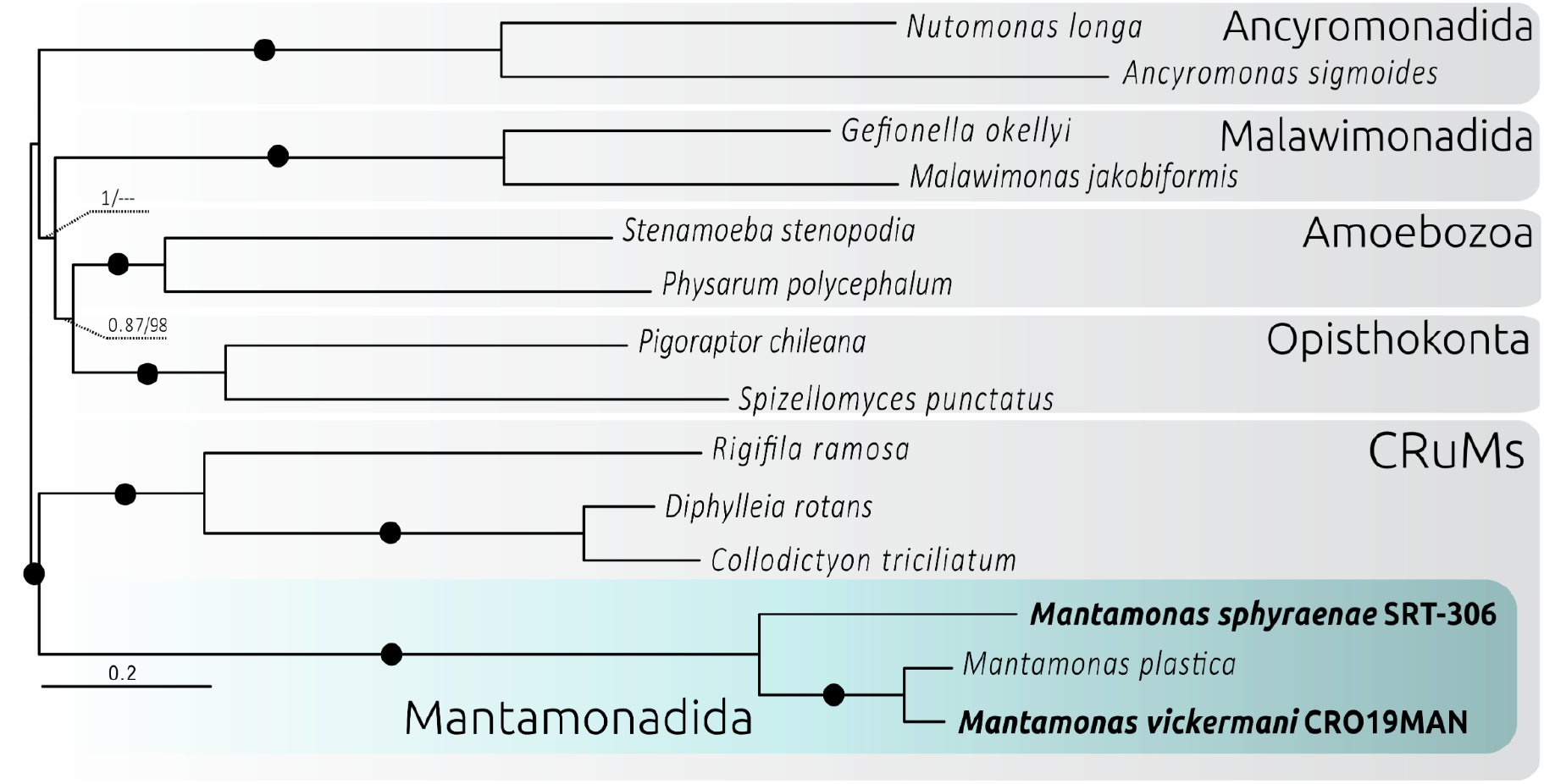
Phylogenomic analysis of CRuMs clade. Bayesian inference (BI) phylogeny based on 182 conserved proteins from Lax et al. (2018). The tree was obtained using 62,088 amino acid positions with the CAT-GTR model. Statistical support at branches was also estimated using maximum likelihood (ML) under the LG+C60+F+R4 model with the PMSF approximation. Numbers at branches indicate BI posterior probabilities and ML bootstrap values, respectively; bootstrap values <50% are indicated by dashes. Branches with support values higher than or equal to 0.99 BI posterior probability and 95% ML bootstrap value are indicated by black dots. The tree was rooted between CRuMs and everything else.

To explore the gene content diversity of our new mantamonad species, we first annotated their genes with EggNOG mapper^9^. 77% and 66% of the predicted proteins from *M. sphyraenae* and *M. vickermani* could be annotated, respectively. A total of 1,718 orthogroups were found to be conserved among all CRuMs taxa (Fig. 5a), while 4,378 were identified as shared between the three *Mantamonas* species, representing the minimal core proteome of the genus *Mantamonas* as currently known (Fig. 5b), among which 2,161 orthogroups are not found in the other two CRuMs lineages.

**Fig. 5.**
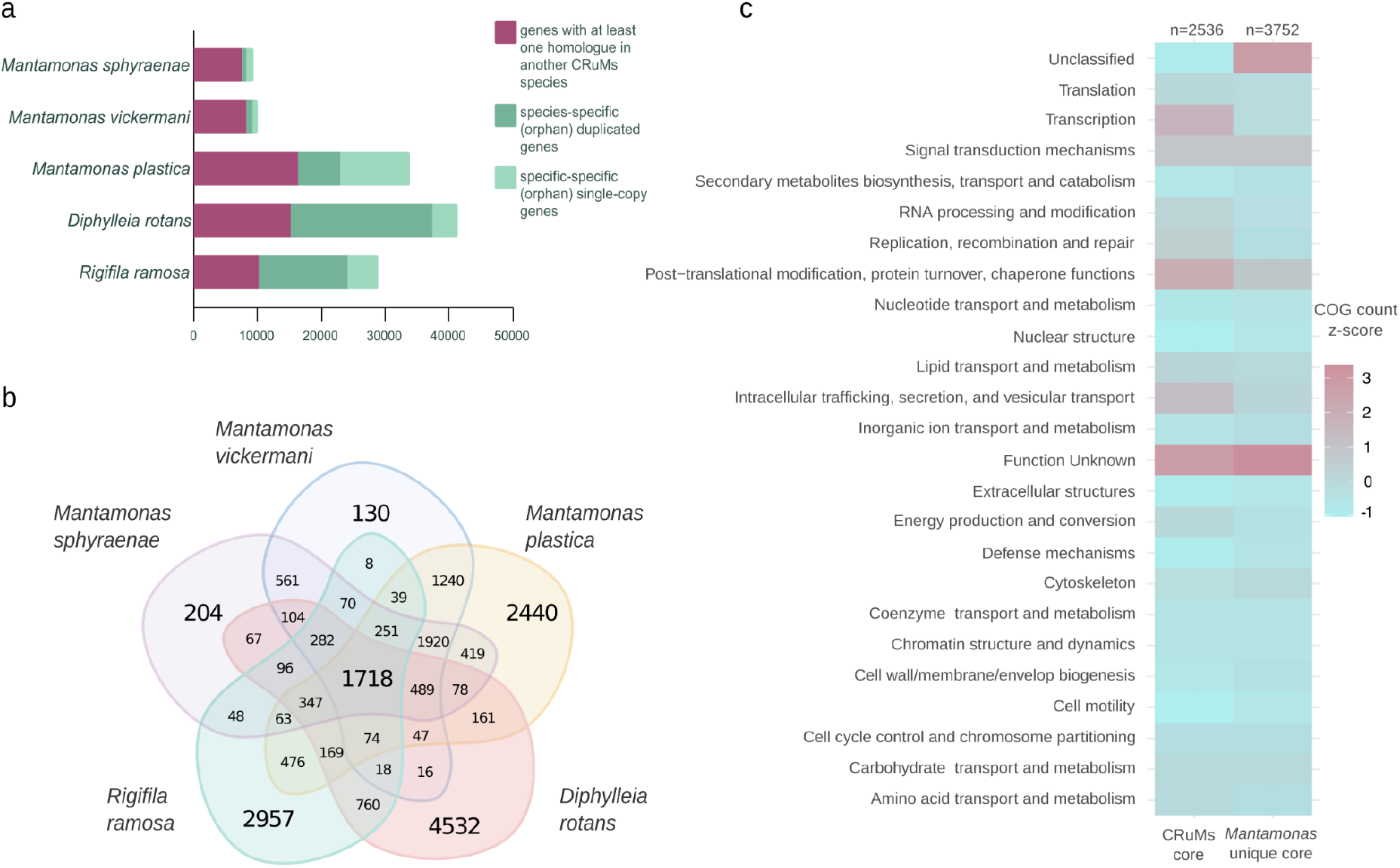
Orthology analysis across the CRuMs supergroup. a) Distribution of genes shared among CRuMs representatives (magenta) or that are species-specific in one or several copies (dark and light green, respectively). b) Number of orthogroups shared among compared CRuMs species. c) COG functional categories associated with orthogroups shared among all CRuMs, as well as those associated with orthogroups shared across *Mantamonas* species but absent in other CRuMs taxa. COG counts were scaled by column using z-score standardization.

Most of the proteins conserved among the CRuMs taxa (99.6%) were found to have an ortholog in the EggNOG database and to belong to at least one Cluster of Orthologous Groups (COG)^10,11^ functional category, where the most highly represented were “Function unknown” and “Post-translational modification, Translation, and Intracellular trafficking” (Fig. 5c). By contrast, a substantial amount of orthogroups conserved among mantamonads (12%), but absent in other CRuMs lineages, could not be assigned to any cluster in the EggNOG database. In addition, most orthogroups conserved in mantamonads but absent in other CRuMs that could be connected to an existing EggNOG cluster were annotated as “Function unknown” (Fig. 5c). Altogether, this large number of *Mantamonas*-specific genes of unknown function suggests that many genetic innovations occurred during the early evolution of this group.

Finally, as an additional way of assessing the *M. sphyraenae* and *M. vickermanii* sequence data completeness and to capture a sense of the complexity of the cellular systems in these organisms, we interrogated the complement of one well-studied set of proteins, the membrane-trafficking system. This complex protein machinery underpins normal cellular function and is critical for feeding, cell growth, and interaction with the extracellular environment^12^. The protein complement associated with the membrane trafficking system has now been investigated in diverse eukaryotic representatives^12^. While some proteins are highly conserved across lineages, others have rarely been retained during evolution but were nonetheless present in the Last Eukaryotic Common Ancestor (LECA)^12^. Among them, the so-called “jotnarlogs” represent LECA proteins present in diverse extant eukaryotes but not in the major opisthokont model organisms.

We detected most proteins of the membrane-trafficking system in the two new *Mantamonas* species, making it one of the most complete known protein complements for this system when compared to representatives of well-characterized model organisms from other supergroups (Fig. 6). Notably, *Mantamonas* encodes some rarely retained proteins, such as the AP5 complex^13^ and syntaxin 17^14^. We also identified several jotnarlogs (Fig. 6), including a near-complete TSET complex, and the SNAREs NPSN and Syp7.

**Fig. 6.**
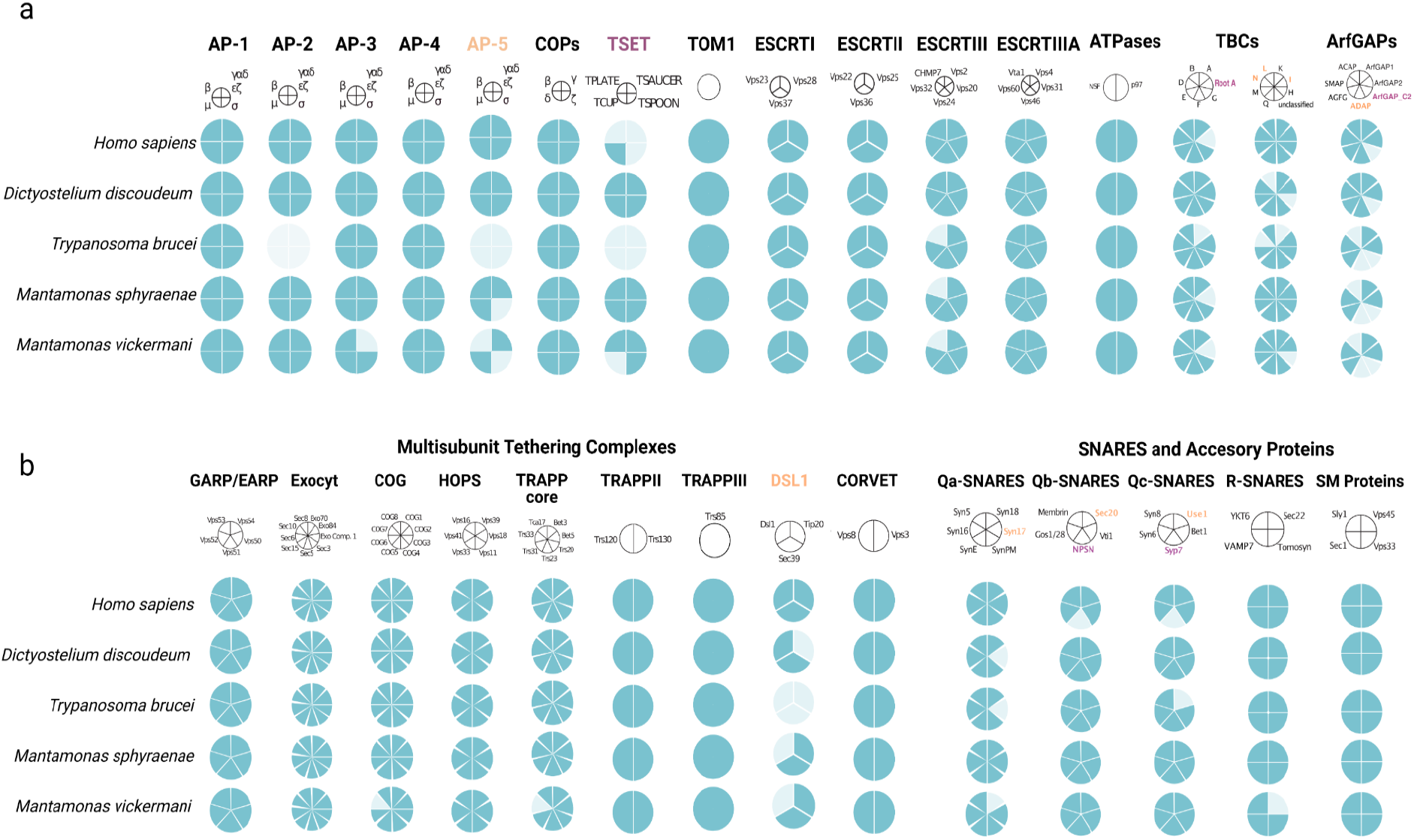
Distribution of proteins associated with the membrane trafficking system in new *Mantamonas* species and other model organisms. a) selected vesicle formation machinery; b) selected vesicle fusion machinery. Names of proteins with jotnarlogs are in purple; those with patchy distribution are in orange.

The identification of homologs of the majority of the protein complement associated with the membrane trafficking system corroborated the high completeness of our genomic and transcriptomic datasets. Interestingly, the presence of proteins previously reported with patchy distributions is consistent with their ancient nature, and, more speculatively, with the hypothesis that free-living flagellates may have retained them from a complex LECA.

Overall, our new *Mantamonas* nuclear genome and transcriptome sequences provide high quality data for a major, yet poorly known, eukaryotic supergroup. They will allow more comprehensive comparative studies of genetic diversity in microbial eukaryotes and a better understanding of deep eukaryotic evolution.

## Methods

### *Mantamonas sphyraenae* isolation, cell culturing, nucleic acid extraction and sequencing

*Mantamonas sphyraenae* SRT-306 was collected on 26 Sep. 2013 from the surface of a barracuda caught in a lagoon on Iriomote Island, Taketomi, Okinawa Prefecture, Japan (24° 23’ 36.762” N, 123° 45’ 22.572” E). It was isolated manually from the rough sample with a micropipette, and maintained in Erd-Schreiber medium^15^ fortified with 2.5% (final volume) freshwater Cerophyl medium (ATCC 802). Stock cultures were kept in 8 ml volumes in 25 ml culture flasks at 16-18°C, and transferred at three-week intervals. Bulk cultures were grown at room temperature in 10 cm Petri plates containing ∼10 ml medium.

To obtain nucleic acids, initially, five plates were inoculated with 500 µl from mature stock cultures. When these had reached high density (qualitatively determined), for each plate, the supernatant was discarded, cells were collected with the use of disposable cell scrapers, and the resulting 0.3–0.5 ml of concentrated cells were inoculated into 50 ml of fresh medium, which was then distributed into five new plates. This process was repeated, for a final count of 125 plates. Cells were harvested with disposable cell scrapers and centrifuged to obtain cell pellets. Independent bulk cultures were used for DNA and RNA extractions.

About 45 µg of total DNA were isolated by using a standard phenol/chloroform/isoamyl alcohol phase-separation protocol and sequenced using Single Molecule Real Time (SMRT) cell technology in a PacBio RSII system at the Cold Spring Harbor Laboratory. A total of 2,304,908 reads (18.7 Gbp) were acquired from 33 SMRT cells. Additional DNA samples were used to prepare two DNA Nextera libraries and a total of 62,929,978 read pairs (18.9 Gbp) and 53,901,870 read pairs (16.2 Gbp) from the short insert and mate-pair libraries, respectively, were generated in an Illumina HiSeq2500 platform at Weill Cornell’s Genome Resources Core Facility. Finally, RNA was isolated using TRI reagent (Sigma) according to the manufacturer’s instructions, using spin columns for elution. The total RNA sample was subjected to poly-A selection followed by Illumina TruSeq RNA library preparation and a total of 24,187,884 read pairs (7.3 Gbp) were sequenced using Illumina HiSeq2500.

### *Mantamonas sphyraenae* genome assembly, gene prediction and ploidy analysis

Both the PacBio and Illumina reads were screened for contamination (see details in the technical validation section) and more than 60% of the original data identified as contaminant was discarded (see technical validation section). After this initial decontamination step, a total of 5.89 Gbp of long read data was assembled using the Canu^5,16^ and FALCON^6^ pipelines.

The resulting genomic contigs from the Canu and FALCON approaches were then polished by aligning the screened PacBio reads to the draft genome using minimap2^17^ and generating a consensus with Racon v1.3.1^18^. Subsequently, a second step of polishing was performed with the high quality Illumina reads by mapping them with bwa-0.7.15^19^ and using Pilon v1.22^20^ to correct for single base errors.

Additionally, MaSuRCA v3.2.6^7^ was used to generate a hybrid assembly using the PacBio as well as the pair-ended and mate-pair Illumina data that were retained after bacterial read filtering.

After these assembly efforts, remaining bacteria contigs were identified by using a combination of homology searches and tetramer frequency-based binning (see details in the technical validation section). From the Canu assembly, a contig corresponding to mtDNA was identified and removed. Clean assemblies were then assessed based on their contiguity and completeness (Supplementary Table 1) and the FALCON assembly was chosen for further analyses. Because of the specific parameter set utilized, our FALCON analysis did not assemble mtDNA due to its much higher sequence coverage compared to that for the nuclear DNA.

A custom library of repetitive elements was generated for the polished and cleaned nuclear genomic sequence by combining the results of RepeatModeler2^21^ and Transposon-PSI (http://transposonpsi.sourceforge.net/) pipelines. The gathered repeat sequences from both analysis were merged and clustered to generate a single consensus and refined repeat library that was further compared against the Dfam database^22^ to classify the repetitive elements using RepeatModeler^21^ refiner and classifier modules. Repetitive elements identified by this procedure were then masked out of the nuclear genome using RepeatMasker^21^ before the prediction of protein-coding genes. Subsequently, the RNA-seq libraries were mapped against the genome sequence with HISAT-2^23^ to generate spliced alignments, and BRAKER2^24^ was employed to predict the nuclear protein coding genes integrating the extrinsic evidence from the RNA-Seq data.

Ploidy was inferred by assessing the distribution of allele frequencies at biallelic single nucleotide polymorphisms (SNPs) visually, and with modeling^25,26^ using nQuire^26^. Briefly, the Nextera Illumina reads were mapped to the final genome assembly with Bowtie2 v2.3.5.1^27^ and the resulting .bam file was used to calculate base frequencies for each biallelic site. These results were denoised using nQuire. The resulting frequencies were plotted in R version 3.3.3^28^. Finally, we ran the nQuire’s Gaussian Mixture Model (GMM) command, which models the distribution of base frequencies at biallelic sites, and uses maximum likelihood to select the most plausible ploidy model.

### *Mantamonas vickermani* isolation, culturing, RNA purification and transcriptome sequencing

*Mantamonas vickermani* CRO19MAN was isolated from a sediment sample collected in July 2014 from the shallow marine lagoon Malo jezero (42°47’05.9”N 17°21’01.3”E) on the island of Mljet (Croatia, Mediterranean Sea). The sample was taken from the upper layer of the sediments in the shore of the lagoon with a sterile 15 ml Falcon tube at a depth of 10 cm below the water surface and stored at -20 ºC. In September 2019, a small amount of sediment was inoculated in a Petri dish with 5 ml of sterile seawater supplemented with 1% YT medium (100 mg yeast extract and 200 mg tryptone in 100 ml distilled water, as in the protocol from the National Institute for Environmental Studies [NIES], Japan). After observation of some mantamonad cells, serial dilution was performed in a multiwell culture plate to further enrich the culture by transferring 250 µl of culture to a well with 1 ml of fresh 1% YT seawater medium and then retransferring the same volume to a new well, repeating the process 5 times for a total of 24 wells. Single mantamonad cells were then isolated from one of the enriched cultures with an Eppendorf PatchManNP2 micromanipulator using a 65 µm VacuTip microcapillary (Eppendorf) and a Leica Dlll3000 B inverted microscope. This cell was inoculated into 1 ml of growth medium and after 48 h incubation we confirmed an established monoculture of *M. vickermani* CRO19MAN.

This new strain was grown for a week in 75 cm^2^ cell culture flasks with ∼10 ml of medium. Fully grown cultures were collected by gently scratching the bottom of the flasks with a cell scraper to resuspend the gliding flagellates and pooled in 50 ml Falcon tubes to be centrifuged at 10ºC for 15 minutes at 15,000 g. Total RNA was extracted from cell pellets with the RNeasy mini Kit (Qiagen), following the manufacturer protocol. Two cDNA Illumina libraries were constructed after polyA mRNA selection, and these were sequenced using the paired-end (2 × 125 bp) method with Illumina HiSeq 2500 Chemistry v4 (Eurofins Genomics, Germany).

### *Mantamonas vickermani* proteome prediction

Reads coming from possible cross-species contamination were removed using CroCo v1.1^29^. The *M. vickermani* transcriptome was then assembled *de novo* using Spades v3.13.1^30^ with the *rna* mode and default parameters specified. Transcripts were then screened to identify remaining contaminants using the Blobtools2^31^ pipeline and homology searches against a custom database (see technical validation section). Predicted proteins were obtained from the clean transcripts using Transdecoder v2 (http:transdecoder.github.io) with default parameters. Subsequently, CD-HIT^32^ clustering was employed (with a threshold of >=90% of identity) to produce a non-redundant data set of proteins for *M. vickermanii*, and to eliminate falsely duplicated proteins stemming from alternatively spliced transcripts.

### Phylogenomic analyses

The dataset of 351 conserved protein markers from Lax et al.^3^ was updated by BLASTP searches^33^ against the inferred proteomes of representatives of other eukaryotic lineages, including the proteomic data for our two new mantamonad strains. Each protein marker was aligned with MAFFT v.7^34^ and trimmed using TrimAl^35^ with the -automated1 option. Alignments were manually inspected and edited with AliView^36^ and Geneious v6.06^37^. Single-protein trees were reconstructed with IQ-TREE v1.6.11^38^ under the corresponding best-fitting model as defined by ModelFinder^39^ implemented in IQ-TREE^38^. Each single-protein tree was manually inspected to discard contaminants and possible cases of horizontal gene transfer or hidden paralogy. At the end of this curation process, we kept a final taxon sampling of 14 species, including members of Ancyromonadida, Malawimonadida, Opisthokonta, and CRuMs (concatenated alignment and supplementary trees are available at https://doi.org/10.6084/m9.figshare.20411172.v1), and 182 protein markers that were present in all mantamonad species (with at least 80% of markers identified in each taxon). All proteins were realigned, trimmed as previously described, and concatenated, creating a final supermatrix with 62,088 amino acids.

A Bayesian inference tree was reconstructed using PhyloBayes-MPI v1.5a^40^ under the CAT-GTR model^41^, with two MCMC chains, and run for 10,000 generations, saving one of every 10 trees. Analyses were stopped once convergence thresholds were reached (i.e. maximum discrepancy <0.1 and minimum effective size >100, calculated using bpcomp). Consensus trees were constructed after a burn-in of 25%. Maximum likelihood (ML) analyses were done with IQ-TREE v1.6.11^38^, first by calculating the ML tree under the LG+F+R4 model, which was used as guide tree for the PMSF approximation^42^ run under the LG+C60+F+R4 model.

### CRuMs orthologue analysis

Orthologous gene families were identified among the predicted proteomes of *Mantamonas sphyraenae, Mantamonas vickermani* and the publicly available proteomes of *Mantamonas plastica, Diphylleia rotans* and *Rigifila ramosa* as obtained from the EukProt v3 database^43^ using OrthoFinder v2.5.4^44^. For this, we used DIAMOND^45^ (“ultra-sensitive” mode, and query cover >= 50%), an inflation value of 1.5, and the MCL clustering algorithm.

### Protein functional annotation

The predicted protein of *M. sphyraenae, M. vickermanii, M. plastica, D. rotans*, and *R. ramosa* were functionally annotated with the EggNOG-mapper pipeline^9^, using DIAMOND ultra-sensitive mode and all domains of life as the target space. During this process, individual sequences composing the CRuMs orthogroups generated by OrthoFinder were assigned a COG functional category. This information was summarized at the orthogroup level by assigning to each orthogroup a single COG category corresponding to the most frequent annotation of its individual sequences, pending that it represented at least 50% of the sequences within the orthogroup.

### Analysis of the conservation of the membrane-trafficking system complement

To assess the complement of the membrane trafficking system encoded in our *Mantamonas* genome and transcriptome datasets, we performed homologous searches of a selection of protein query sequences from the genomes of *Homo sapiens, Dictyostelium discoideum, Arabidopsis thaliana* and *Trypanosoma brucei* available at the NCBI (National Center for Biotechnology Information) database (Supplementary Table 3). These proteins included components of the machinery for vesicle formation (HTAC-derived coats, ESCRTs, and ArfGAPs) and vesicle fusion (SNAREs and SM proteins, TBC-Rab GAPs, and Multi-subunit tethering complexes)^12^.

BLASTP and TBLASTN were used to search the predicted proteomes and nucleotide coding sequences, respectively, of *M. sphyraenae* and *M. vickermani*. The HMMER3 package was used to find more divergent protein sequences using the hmmsearch tool^46^. In cases in which only TBLASTN hits were retrieved, these were translated using Exonerate^47^. Potential orthologs (i.e., hits with an E-value below 0.05) were further analyzed by the Reciprocal Best Hit (RBH) approach, using the *Mantamonas* candidate orthologs as queries against the *H. sapiens, D. discoideum* and *A. thaliana* proteomes. If the best hit was the protein of interest and had an E-value two orders of magnitude lower than the next non-orthologous hit, this was considered as orthology validation. Forward and reverse searches were performed using the AMOEBAE tool^48^.

### Data records

The sequence data associated with the nuclear genome and transcriptome of *Mantamonas sphyraenae* as well as the transcriptome of *Mantamonas vickermani* (Table 2) have been deposited to the NCBI under the Bioproject accession PRJNA886733.

## Technical validation

### Quality assessment of sequencing datasets

All Illumina paired-end raw reads used for genome polishing were quality-checked with FastQC v0.11.8^49^ and trimmed using TRIMMOMATIC^50^ to retain only reads with maximum quality scores. PacBio reads resulted in an N50 of 11,048 bp and an average coverage of 106x after filtering out the identified contaminant sequences (see below).

### Identification and filtering of contaminant sequences

Mantamonads grow in non-axenic cultures with co-cultured prokaryotes on which they feed. Therefore, various methods were employed to ensure the correct identification and filtering of contaminant sequences of the genomic and transcriptomic datasets of *M. sphyraenae* and *M. vickermani*.

For the genomic dataset of *M. sphyraenae*, we first identified the main bacterial contaminants from the initial genome assemblies ^51^. In addition, we established a customs database consisting of contigs assembled from Illumina sequencing data from bacteria only enrichment cultures derived from the lab’s several xenic protist cultures. These were used to screen PacBio reads using BLASR v5.1^52^ and Illumina reads using Bowtie2 v2.3.5.1^27^. Only Illumina reads in which reads from neither pair aligned to the bacterial database were retained for further assembly.

After genome assembly using the filtered reads, remaining contaminant contigs were identified by using MyCC v1^53^, which bins contigs based on their tetranucleotide frequencies and coverage. Clusters were formed using the affinity propagation (AP) algorithm and visualized in a 2-dimensional Barnes-Hut-SNE plot. BLASTN searches using default parameters were conducted against the ‘nt’ database from the NCBI to taxonomically classify the bins. Contigs were identified as contaminants if they contained no hits other than to prokaryotes, and were clustered away from the main eukaryotic bin. Finally mitochondrial sequences were screened out from the short and long read libraries of *M. sphyraenae* by mapping them against the mitochondrial genome using bwa-0.7.15^19^ and minimap2^17^ respectively.

The assembled transcriptome of *M. vickermani* was decontaminated with the Blobtools2 pipeline^31^. Briefly, this approach helps to identify contaminant sequences based on their biases in coverage and GC content, as well as on a taxonomic classification established by DIAMOND searches^45^ against the ‘nt’ and Uniprot databases^54^. In addition, a second cleaning step was done by performing DIAMOND searches against a database containing all the proteins of the prokaryotic Genome Taxonomy Database (GTDB)^55^ and the eukaryotic-representative EukProt v3 database^43^. A protein was considered as a probable contaminant and excluded from further analyses if its best hit corresponded to any protein from GTDB, with strict cutoffs of identity ≥ 50% and query coverage ≥ 50%. Finally, a blobplot was generated for the final genomic and transcriptomic contigs of *M. sphyraenae* and *M. vickermani*, respectively, to verify the absence of contaminant sequences (Supplementary Figs. 1 and 2).

### Completeness analysis

To assess the completeness of the decontaminated genome and transcriptome datasets, we employed the BUSCO v5.3.2 pipeline. We identified the percentage of near-universal single copy orthologs of the eukaryote_odb10 database^16^ on the predicted proteomes of *M. sphyraenae* and *M. vickermani*, as well as those of other species belonging to the CRuMs supergroup available in the EukProt v3^43^ database for comparison purposes.

## Code availability

All the employed software as well as their versions and parameters were described in the method section. If no parameters were specified, default settings were employed. Data visualization plots were generated using R v4.1.2 (https://cran.r-project.org/, R development core team) and https://bioinformatics.psb.ugent.be/webtools/Venn/.

## Acknowledgements

The authors thank Drs. J.Z. Xiang, D. Xu, and H. Shang at the Weill Cornell’s Genome Resources Core Facility for their assistance with Illumina sequencing. They also thank Dr. S. Goodwin in the NGS sequencing core at Cold Spring Harbor Laboratory for her help with PacBio sequencing, and Drs. J.A. Burns and A.A. Pittis for their initial data analysis efforts. This work was funded by the Simons Foundation Grant awards to EK (SF-382790 & SF-876199). This project has received funding from the European Research Council (ERC) under the European Union’s Horizon 2020 research and innovation programme (ERC Starting Grant No 803151 to L.E., and ERC Advanced Grants No 322669 and 787904 to P.L.-G. and D.M., respectively). L.J.G. was funded by the Horizon 2020 research and innovation programme under the European Marie Skłodowska-Curie Individual Fellowship H2020-MSCA-IF-2020 (grant agreement no. 101022101 - FungEye). GT was supported by the 2019 BP 00208 Beatriu de Pinos-3 Postdoctoral Program (BP3; 801370). Work in the Dacks lab is supported by Discovery Grants from the Natural Sciences and Engineering Research Council of Canada (RES0043758, RES0046091).

## Author contributions

JB: Conceived and designed the analysis; Contributed data or analysis tools; Performed the analysis; Wrote the paper.

LJG: Collected the data; Performed the analysis.

AA: Collected the data; Performed the analysis; Wrote the paper.

HK: Performed the analysis.

GT: Contributed data or analysis tools; Performed the analysis.

AY: Conceived and designed the analysis; Collected the data; Contributed data or analysis tools; Performed the analysis; Wrote the paper.

LAT: Performed the analysis. AF: Performed the analysis.

SW: Conceived and designed the analysis; Performed the analysis.

AP: Contributed data or analysis tools; Performed the analysis.

TS: Collected the data.

KI: Contributed data or analysis tools.

JBD: Conceived and designed the analysis; Contributed data or analysis tools; Wrote the paper.

PLG: Contributed data or analysis tools.

DM: Contributed data or analysis tools; Wrote the paper.

EK: Conceived and designed the analysis; Contributed data or analysis tools; Wrote the paper.

LE: Conceived and designed the analysis; Contributed data or analysis tools; Wrote the paper.

## Supplementary Information

**Taxonomy section (microscopy and formal species description)**

**Table S1. Genome assembly statistics**

**Table S2. nQuire Gaussian Mixture Model delta log-likelihood values**

**Table S3. Membrane-trafficking system complement**

**Fig. S1. *M. sphyraenae* Blob plot**

**Fig. S2. *M. vickermani* Blob plot**

**Fig. S3. Bayesian phylogenomic tree**

**Fig. S4. Maximum Likelihood phylogenomic tree**

