## Supplementary Information for "High quality genome and transcriptome data for two new species of *Mantamonas*, a deep-branching eukaryote clade"

### Taxonomy

Microscopy and morphological description of *Mantamonas vickermani* sp. nov.

Optical microscopy observations were performed with a Zeiss Axioplan 2 microscope equipped with oil-immersion differential interference contrast (DIC) and phase contrast objectives. Images were acquired with an AxiocamMR camera using the Zeiss AxioVision 4.8.2 SP1 suite. Videos were recorded using a Sony α9 digital camera (Movie S1 and Movie S2). Morphometric data were obtained at 1,000x final magnification on 20 cells. Images were captured at multiple focal planes in order to visualise different cell parts. Measurements of flagella pertain to the visible parts, i.e., the posterior flagellar length is measured beginning from the point at which it emerges from underneath the cell at the body's posterior end.

*Mantamonas vickermani* cells are ~3 µm wide and ~3.5 µm long; thus noticeably smaller than those of *Mantamonas plastica* (~5 µm wide and ~5 µm long) (Glücksman et al. 2011) (Figure 1g-l). Like *M. plastica*, *M. vickermani* also has a strongly flattened and plastic morphology. However, the characteristic blunt projection on the left-hand side of the cell observed in *M. plastica* is less conspicuous in *M. vickermani*, and not always observed in cells possessing an overall spherical to oval morphology (Figure 1). The anterior flagellum of *M. vickermani* is ~2 µm long, rigid in all of its length, and vibrates with a small amplitude; its posterior flagellum is ~7 µm long and considerably thicker than the anterior one, having a very small acroneme that when seen is never longer than 1-2 µm.

Both flagella are also shorter than those reported for *M. plastica* (~3 µm anterior and ~ 10 µm posterior).

*Mantamonas vickermani* glides in a smooth and continuous manner on the substrate with a similar speed and turning behavior to that observed for *M. plastica* (Glücksman et al 2011; AAH, pers. obs.) (Movie S1-S2). As with *M. plastica*, *M. vickermani* is a bacterivore with a voracious appetite, engulfing bacteria at a high rate. Interestingly, and in contrast with Glücksman et al. (2011), we did observe one cell possessing two posterior flagella, which strongly suggests that it was undergoing cellular division (Figure 1l).

<u>Microscopy and morphological description of *Mantamonas sphyraenae* sp. nov.</u>

Live cells were observed on an Zeiss Axiovert 100M inverted microscope equipped with DIC and phase contrast optics. Images were captured with an Olympus DP73 17.28-megapixel camera. As for *M. vickermani*, morphometric data were obtained at 1,000x magnification on 20 cells. *Mantamonas sphyraenae* cells exhibited three general morphologies: 'balloons', which were typically ~5 µm long and ~3 µm wide, with a circularly curved anterior and a posterior end tapering to a point; 'kites', which were roughly diamond-shaped, about 3.5–4 µm long and wide; and 'mantas', which were 4–5 µm wide and ~3 µm long, having a broadly curved anterior end, a more tightly rounded right side, a bluntly rounded projection on the left side, and a posterior comprising either straight edges culminating in a point, two shallowly concave curves, or one of each. All three morphologies were plastic to some extent, although 'mantas' were noteworthy in that the left-side projection appeared rigid, and the curved right side frequently very plastic. Intermediates between the three morphologies were sometimes observed. In general, all cells in any given culture flask exhibited the same morphology, which often changed from one observation to the next, one to three weeks later. Exceptions to the prevalent morphology were almost always intermediate forms. We did not observe active transitions from one cell type to another, including to or from intermediate forms. Cells of all morphologies glided slowly and with constant speed, although occasionally stopping; the cell body frequently deformed when changing direction or colliding with other objects.

In all cases, a flagellum, 6–10 µm long, trailed behind the cell, always in a straight line except when the cell was turning, in which case it followed the cell's path. No movement of the flagellum was seen besides this. Under extremely favourable conditions, a second flagellum could be seen projecting from the anterior-left of 'manta' cells, at about a 45° angle. This second flagellum was invariably very thin, stiff, and 1–2 µm long. Very occasionally, we observed an additional protrusion, about the full width of a flagellum and about 1–2 µm long. This was always seen projecting from the posterior of the cell, immediately to the left of, and usually parallel to, the posterior flagellum. It appeared entirely static, and never appeared to change its length or orientation. Cysts were never observed at any stage of culture. Likewise, we never observed dividing cells.

**Formal species descriptions**

All taxonomic descriptions in this work were approved by all authors.

Eukarya: 'CRuMs'
Order Mantamonadida Cavalier-Smith 2011
Family Mantamonadidae Cavalier-Smith 2011
Genus *Mantamonas* Cavalier-Smith and Glücksman 2011
*Mantamonas sphyraenae* sp. nov.

**Description:** Cells with varying morphologies: shaped as manta rays (as for genus in Glücksman et al. 2011), ~3 µm long and ~5 µm wide; diamonds, 4±1 µm in both dimensions; or rounded anteriorly and tapering posteriorly, ~5 µm long and ~3 µm wide. Anterior flagellum stiff, 0.5–1.0 µm long. Other characters as for genus.
**Type culture:** SRT306
**Type locality:** Surface of barracuda caught in lagoon on Iriomote Island, Taketomi, Okinawa Prefecture, Japan (24° 23' 36.762" N, 123° 45' 22.572" E).
**Isolator:** Takashi Shiratori
**Etymology:** From *Sphyraena*, the genus name for barracuda, the fish from which the type strain was obtained.
**Gene sequence:** The nuclear genome and transcriptomic read sequencing data from *Mantamonas sphyraenae* (strain SRT306) were deposited in GenBank under BioProject accession number PRJNA886733.

*Mantamonas vickermani* sp. nov.

**Description.** Cell size ~3 µm (2.5–4.3 µm) long, ~3.5 µm (3.0–4.0 µm) wide; cells almost perfectly round, although in some cases possessing a small projection to the left side of the cell; without pseudopodia; anterior flagellum usually ≤2 µm long (1.2–2.7 µm), held forwards and to left ~40–50° to longitudinal axis, does not beat except for slight terminal vibration; posterior flagellum ~7 µm long (6–8.9 µm), conspicuous and sometimes acronematic. Other characters as for genus.

**Type culture.** CRO19MAN

**Type locality.** Specimen isolated from the sediments of the marine lake Malo jezero in the island of Mljet, Croatia.

**Isolator:** Luis Javier Galindo.

**Etymology.** The name *vickermani* honors work on heterotrophic protists by Keith Vickerman.

**Gene sequence.** The full transcriptome read data from *Mantamonas vickermani* (strain CRO19MAN) were deposited in GenBank under BioProject accession number PRJNA886733.

### Supplementary Tables.

Table S1. *Mantamonas sphyraenae* genome assembly statistics

| Assembly approach | Canu | Falcon | MaSuRCa |
|---|---|---|---|
| Total length | 27350505 | 25061354 | 26111353 |
| Number of contigs | 172 | 78 | 136 |
| Mean length | 159014.56 | 321299.41 | 191995.24 |
| Longest contig | 732584 | 751365 | 1133621 |
| Shortest contig | 20756 | 17266 | 1222 |
| N_count | 0 | 0 | 4688 |
| Gaps | 0 | 0 | 9 |
| N50 | 303774 | 375077 | 386663 |
| N50n | 32 | 26 | 24 |
| N70 | 224361 | 300753 | 269297 |
| N70n | 52 | 41 | 40 |
| N90 | 50673 | 226430 | 146039 |
| N90n | 95 | 60 | 65 |
| BUSCO eukaryota odb10 | C:89.1%[S:82.0%,D:7.1%], F:2.0%,M:8.9% | C:91.4%[S:90.6%,D:0.8%], F:2.0%,M:6.6% | C:89.8%[S:86.7%,D:3.1%], F:2.0%,M:8.2% |

Table S2. nQuire Gaussian Mixture Model delta log-likelihood values for the *Mantamonas sphyraenae* genome

| Genome ploidy | Delta log-likelihood values |
|---|---|
| Diploid | 10,944 |
| Triploid | 161,437 |
| Tetraploid | 104,902 |

### Supplementary Figures.

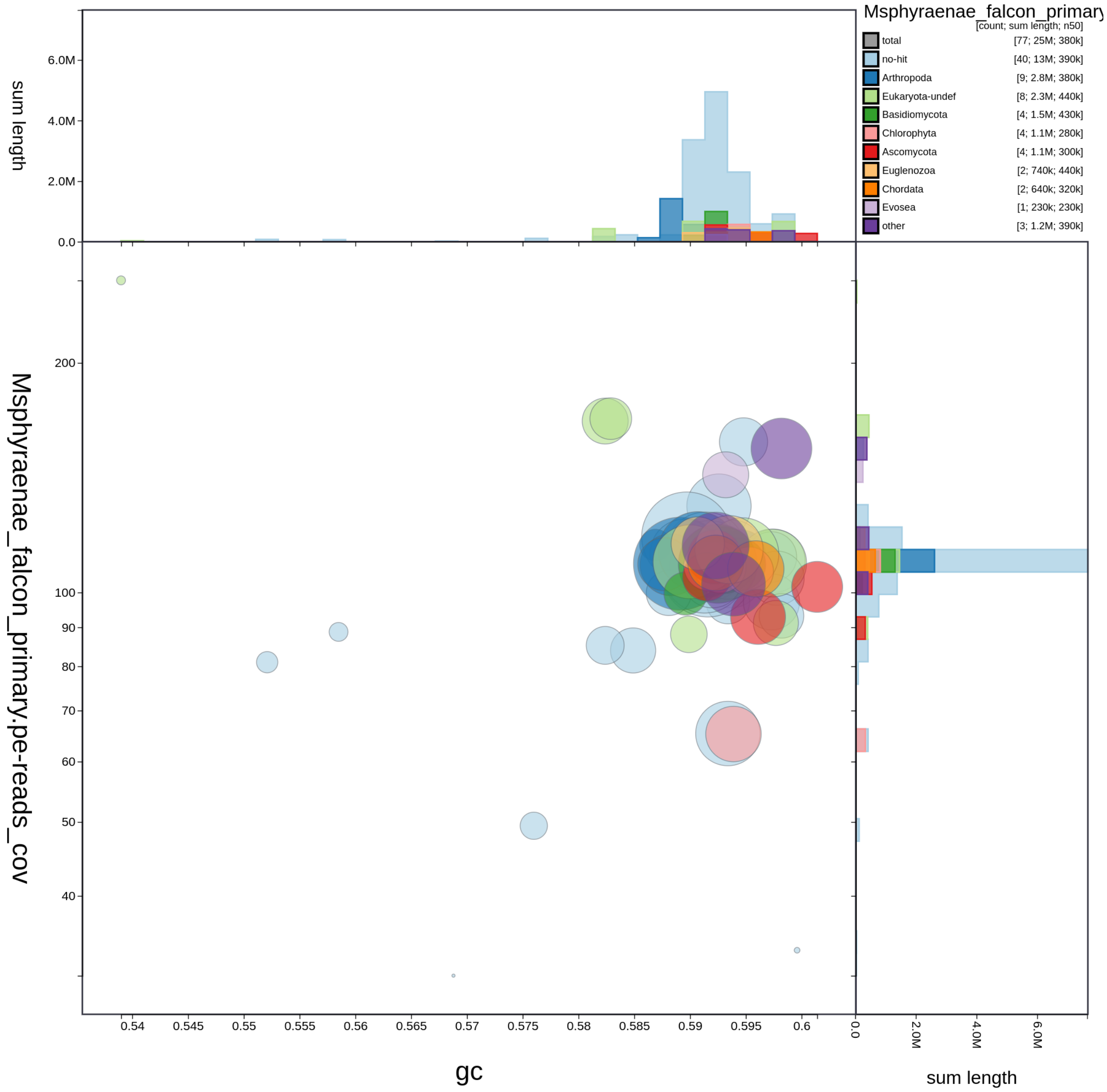


**Supplementary Figure 1.** Blob plot of read coverage against GC proportion in *M. sphyraenae* genomic contigs. Records are coloured according to their similarity to different phyla. Circles are sized in proportion to records cumulative length. The assembly has been filtered to exclude records whose taxonomic assignment matches "Bacteria". Histograms show the distribution of record length sums along each axis.

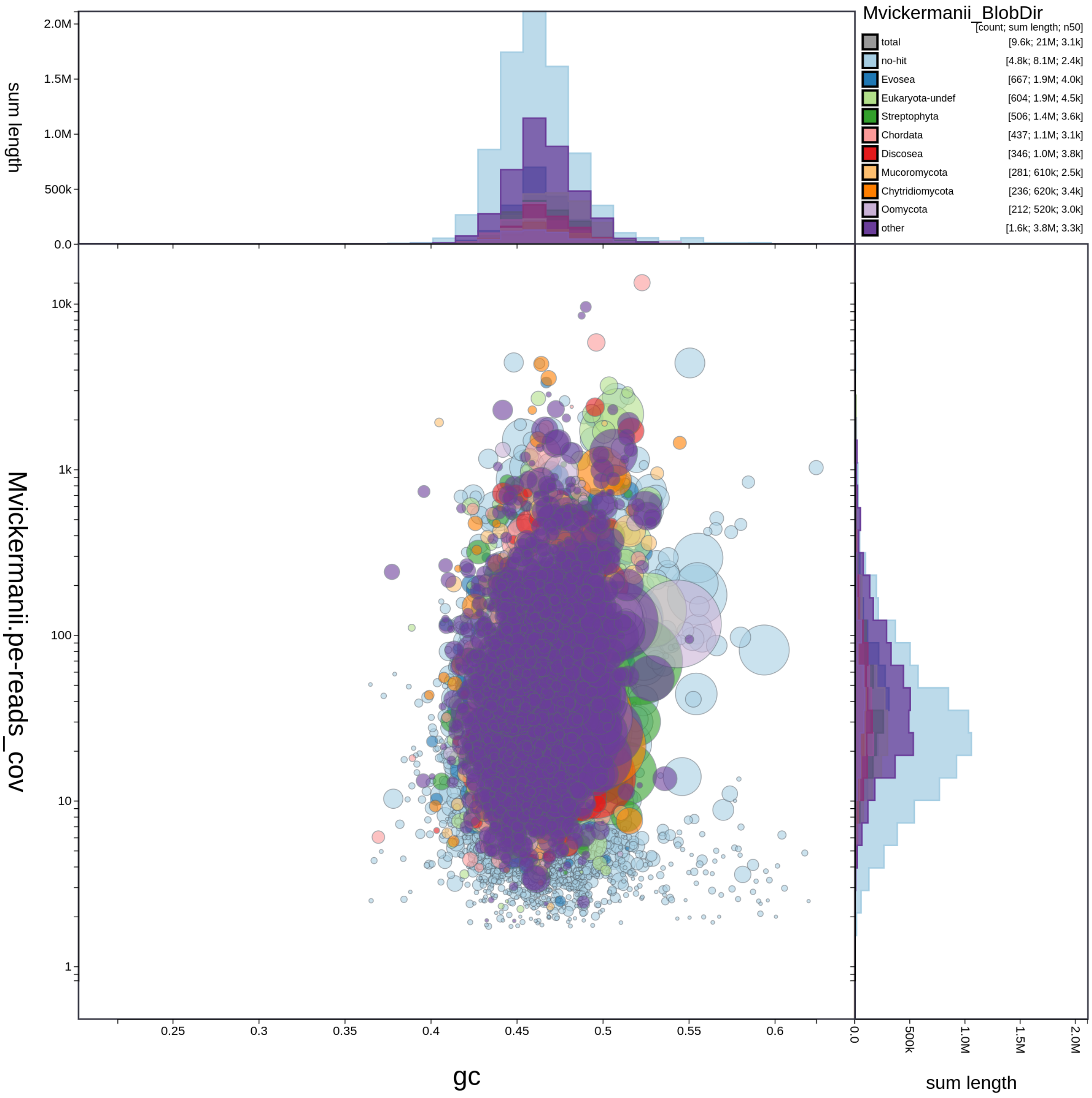


**Supplementary Figure 2.** Blob plot of read coverage against GC proportion in *M. vickermanii* transcriptomic contigs. Records are coloured according to their similarity to different phyla. Circles are sized in proportion to records cumulative length. The assembly has been filtered to exclude records whose taxonomic assignment matches "Bacteria". Histograms show the distribution of record length sums along each axis.

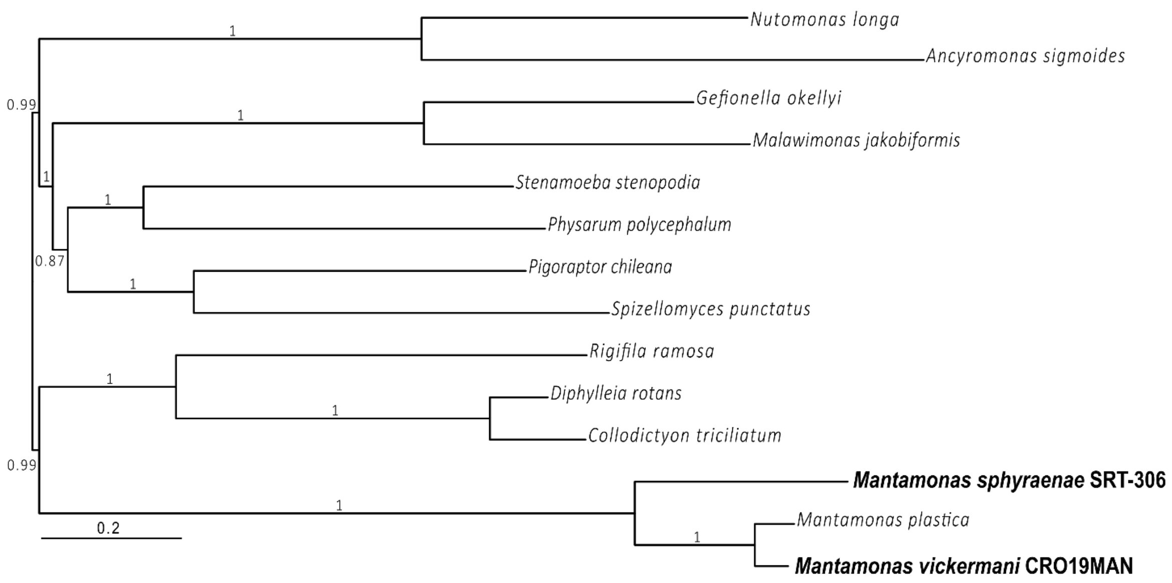


**Supplementary Figure 3.** Bayesian phylogenomic tree based on an updated dataset from Lax et al. 2018. The tree was reconstructed using 182 conserved proteins, 14 taxa, and 62,088 conserved amino acid positions. It was inferred using PhyloBayes under the CAT-GTR model, with posterior probability as branch statistical support.

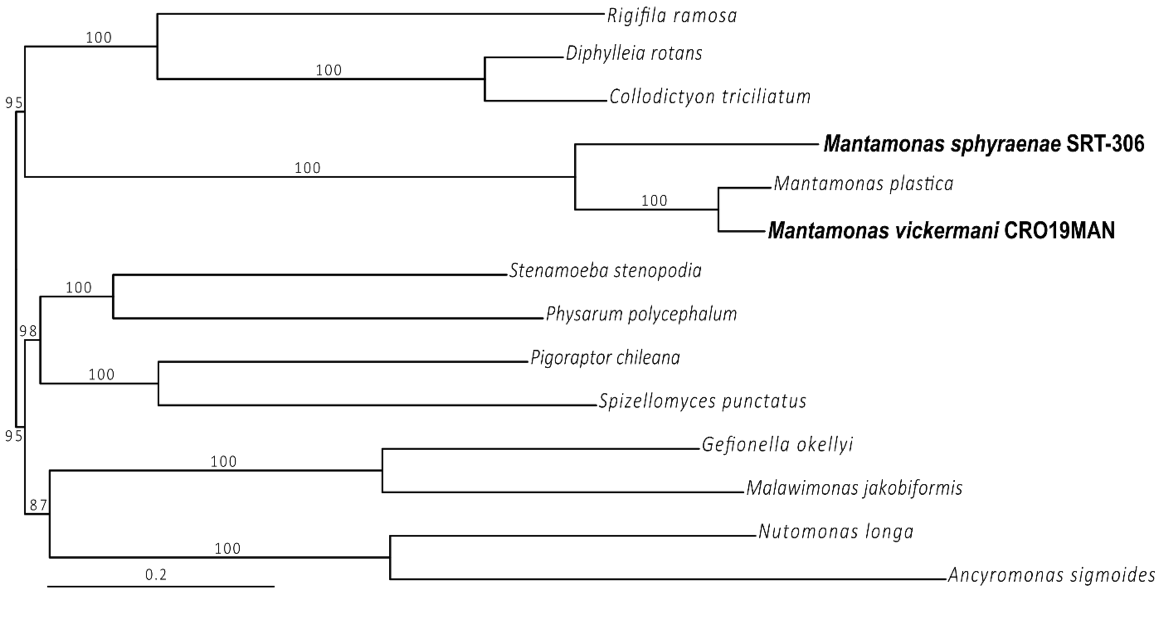


**Supplementary Figure 4.** Maximum likelihood tree based on updated dataset from Lax et al. 2018. The tree was reconstructed using 182 conserved proteins, 14 taxa, and 62,088 conserved amino acid positions. It was inferred with IQ-TREE using the PMSF approximation of the LG+C60+F+R4 model and ultrafast bootstrap as branch statistical support.
